# CRISPR-Engineered hiPSC-derived Cardiomyocytes Reveal Divergent Responses to Loss and Defective Processing of A-type Lamins

**DOI:** 10.64898/2026.06.29.735225

**Authors:** Lauran Vandeweyer, Elisa Garrido-Huéscar, Mirthe Vandenputte, Bert Vandendriessche, Maaike Alaerts, Laura Ordovás, Bart Loeys, Winnok H. De Vos

## Abstract

A-type lamins are critical for nuclear integrity and mechanotransduction in cardiomyocytes, and their disruption is a major cause of inherited cardiomyopathy. To compare the consequences of lamin A/C loss versus defective lamin A maturation, we generated CRISPR/Cas9-edited hiPSC lines lacking *LMNA* or *ZMPSTE24* and differentiated them into iPSC-derived cardiomyocytes. *LMNA* knockout caused progressive nuclear deformation, loss of culture stability, and contractile vulnerability in iCM. *ZMPSTE24* knockout led to subtler nuclear abnormalities and reduced calcium transient activity, temporally correlating with prelamin A accrual. Transcriptomics profiling revealed aberrant mechanical responses in both *LMNA* and *ZMPSTE24* bi-allelic knockouts as well as unique perturbations in inflammatory signaling and epigenetic pathways. Interestingly, both knockout models shared a marked defect in proteostasis, as confirmed by reduced proteasome activity. Together, these results show that loss of lamin A/C and accumulation of prelamin A trigger both converging and distinct cardiomyocyte stress responses. In addition, the newly generated models offer an attractive platform to study lamin-associated cardiomyopathy and its therapeutic targeting.

## Introduction

Cardiomyopathies are disorders that affect the heart muscle function and compromise its elementary function of pumping blood through the body. Dilated cardiomyopathy (DCM) is the most common subtype, with an estimated prevalence of 40 cases per 100 000 individuals worldwide. It is characterized by dilation of the left or both ventricles and typically presents with electrical abnormalities and arrhythmia, which culminates into heart failure. With no curative treatment available, it is the leading cause of heart transplants (1,2). DCM can have an acquired or genetic (hereditary) origin, with at least one third having an autosomal dominant inheritance pattern, usually presenting in the second or third decade of life (3,4). Mutations in the *LMNA gene* are the most commonly detected genetic cause of familial DCM, but the mutational landscape is diverse and genotype - phenotype correlation are poorly understood (5). *LMNA* encodes the A-type lamins, including alternative splice variants lamin A and lamin C, that, together with B-type lamins form the nuclear lamina, a filamentous network of intermediate filaments at the intranuclear face of the nuclear envelope (NE). While B-type lamins are expressed from the earliest developmental stages in all cell types, expression of A-type lamins is restricted to specific cell types and initiates in the later phases of embryonic development (6). Lamin A is produced as a precursor protein, prelamin A, which undergoes significant post-translational modifications, unlike lamin C, which is translated directly as a mature protein. First, prelamin A is farnesylated on the cysteine residue of the terminal CAAX motif, followed by a proteolytic removal of the three terminal amino acids by the zinc metallopeptidase STE24 (ZMPSTE24) or RCE1, and a methylation of the new terminal cysteine. Prelamin A then undergoes a proteolytic reaction in which the last 15 amino acids, including the farnesyl tail, are cut off by ZMPSTE24. The lamina provides structural support for the nucleus and integrates mechanical cues from the environment, through linker of nucleo- and cytoskeleton (LINC) complexes that span the NE. As a consequence, diverse mutations in the *LMNA* gene have been shown to render nuclei from different cell types more susceptible to mechanical stress (7). A balance between the cytoskeletal forces and nuclear resistance (coupled through the LINC complex) is essential for cell homeostasis (8). This is particularly important for cardiomyocytes (CM) as they endure extreme mechanical stress from contraction - relaxation cycles (9). In line with this, disruption of the LINC complex ameliorates *LMNA*-associated nuclear damage in CM by eliminating the perinuclear microtubule cage encircling the nucleus (10,11). However, not all aspects of *LMNA*-associated DCM pathogenesis can be attributed to structural defects. For example, a mouse model with a cardiomyocyte specific Lamin A/C depletion revealed Golgi and endoplasmic reticulum (ER) stress via activation of the ATF4-CHOP axis (12). Mice expressing mutant *LMNA* (p.Asp300Asn), a mutation associated with progeroid syndromes, showed pathogenic activation of E2F/DNA damage response/TP53 pathway in the heart ultimately leading to myocardial fibrosis, apoptosis, cardiac dysfunction and premature death (13). And, myopathic mutations perturbing the tertiary structure of the A-type lamin Ig-fold domain had minimal impact on the nuclear stiffness, but upregulation of Nrf2 target genes indicative of reductive stress (14). Furthermore, several *LMNA* mutations do not necessarily reduce lamin A/C levels but instead exert pathogenic effects through altered protein structure or processing. For instance, missense mutations in the rod domain, such as p.Glu161Lys (E161K) and p.Arg190Pro (R190P), are associated with DCM and primarily disrupt lamin assembly and nuclear mechanics (15,16). In contrast, mutations near the C-terminal processing region, such as p.Leu647Arg (L647R) and p.Arg644Cys (R644C), impair lamin A maturation by reducing ZMPSTE24-mediated cleavage efficiency, leading to accumulation of farnesylated prelamin A. These mutations are typically associated with more variable, often multisystem laminopathy phenotypes that can include cardiac involvement (17–20). Collectively, these observations indicate that *LMNA*-associated cardiomyopathy can arise through at least two fundamentally distinct, yet potentially overlapping mechanisms: loss of functional lamin A/C and accumulation of dysfunctional lamin A species, including prelamin A. Despite increasing recognition of this mechanistic heterogeneity, their relative contributions to cardiomyocyte dysfunction remain poorly defined, particularly in human systems. To dissect these mechanisms in a controlled and genetically defined context, we generated CRISPR/Cas9-edited hiPSC lines carrying mono- and biallelic null mutations in *LMNA* or *ZMPSTE24*, and differentiated them into CM. This approach allowed us to separate the effects of lamin A/C loss from those of dysfunctional lamin A accumulation within a controlled context. Through integrated morpho-functional and transcriptomic analyses, we identified both shared and distinct stress responses, including a common defect in proteasome activity that may contribute to cardiomyocyte dysfunction.

## Results

### CRISPR/Cas9-mediated genome editing establishes stable knockout hiPSC clones for LMNA and ZMPSTE24

To generate stable *LMNA* and *ZMPSTE24* knockout hiPSC lines, we used CRISPR/Cas9 targeting exon 1 of each gene with specific guide RNAs (Fig. 1A,B). Individual clones were obtained after three rounds of low-density seeding followed by single-colony picking, and editing of the target locus was confirmed by Sanger sequencing. All selected CRISPR-edited hiPSC clones retained stemness, as shown by a comparable percentage of OCT4-positive nuclei relative to the parental line (Fig. S1A-B). In addition, a trilineage assay confirmed preservation of differentiation capacity, based on the induction of ectodermal, mesodermal and endodermal-marker transcripts (MAP2, *NKX2.5* and *SOX17*, respectively) (Fig. 1C).

**Figure 1.**
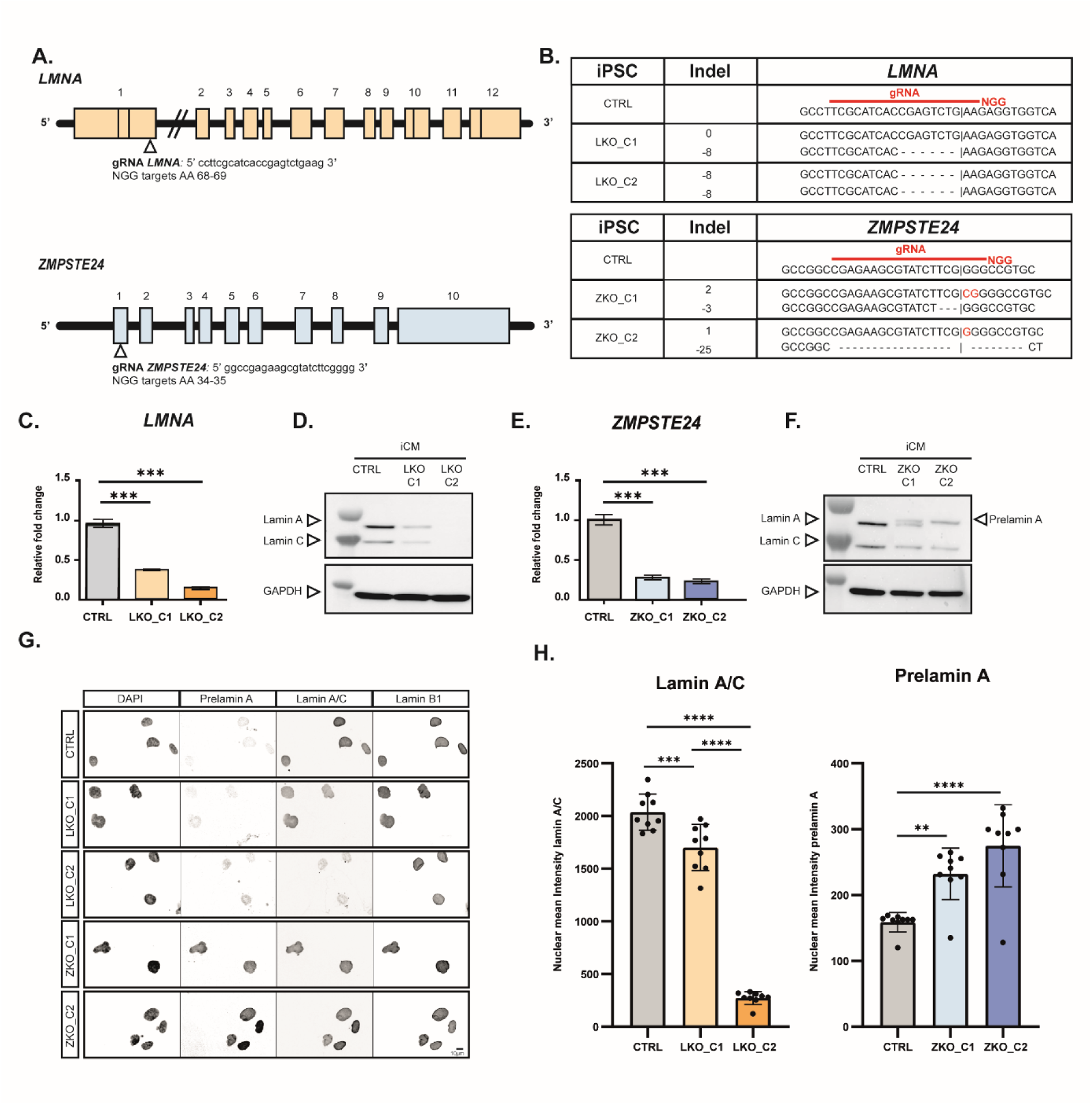
Creation of stable *LMNA* and *ZMPSTE24* knockout hiPSC clones. (A) Schematic of *LMNA* and *ZMPSTE24* gene. The region of the NGG-target site is indicated for the two designed sgRNA. (B) Sanger sequencing to validate CRISPR after three rounds of colony selection on hiPSC level. Two colonies are selected for each targeted gene (*LMNA* and *ZMPSTE24*). (C) qPCR analysis for *LMNA* exon 1 performed on the D12 iCM (n=3, Mann Whitney test, p<0.05) (D) Western blot analysis on CTRL and LKO_C1 & C2 iCM D12 samples for lamin A/C. *GAPDH* is used as reference gene. (E) qPCR analysis for *ZMPSTE24* exon 1 performed on the D12 iCM (n=3, Mann Whitney test, p<0.05) (F) Western blot analysis on CTRL and ZKO_C1 & C2 iCM D12 samples for lamin A/C. (G) Montage of representative images of immunofluorescence staining on D12 iCM for DAPI, prelamin A, lamin A/C and lamin B1. Scale bar = 10µm. (H) Quantification of mean intensity immunofluorescence staining after segmentation nuclei using Cellblocks. (n_bio_=3, n_tech_=3, linear mixed model).

For *LMNA,* two clonal lines were selected: LKO_C1, carrying a heterozygous 8-nucleotide deletion (c.192_199del, p.Thr64-Glu67del), and LKO_C2, carrying a homozygous 8-nucleotide deletion introducing a frameshift (c.192_199del; p.Thr64-Glu67del / (c.192_199del, p.Thr64-Glu67del). These clones were further characterized by qPCR, western blotting (WB) and immunofluorescence (IF) (Fig. 1C-H). Because *LMNA* expression is only initiated upon differentiation, knockout efficiency was assessed in early iCM at D12 (Fig. S1D-G). qPCR using primers spanning the target region revealed strongly reduced transcript levels in both clones compared with the parental control line (CTRL) (Fig. 1C). At the protein level, LKO_C2 showed no detectable lamin A/C signal by WB and only non-specific, punctate staining in IF images, while LKO_C1 showed reduced lamin A/C by WB and a decreased nuclear IF intensity, consistent with partial expression loss (Fig. 1D, G & H).

For *ZMPSTE24*, Sanger sequencing identified one compound heterozygous clone, ZKO_C1, with a 2-nucleotide insertion in one allele and a 3-nucleotide deletion in the other allele (c.61_63insCG; p.Gly21AlaFs / c.del59_61; p.20_21delins), and one compound heterozygous clone, ZKO_C2, carrying a 1-nucleotide indel in one allele and a 25-nucleotide indel in the other allele (c.61_62insC; p.Ala22GlyFs / c.del45_69; p.Ala15_Val23del) (Fig. 1B). Both clones exhibited significantly reduced *ZMPSTE24* transcript levels in D12 iCM. In addition, both IF and WB demonstrated accumulation of prelamin A. On WB, prelamin A accumulation was visible as an additional higher band in ZKO_C1 and a complete band shift in ZKO_C2, relative to the lamin A band, consistent with partial resp. complete impairment of prelamin A processing (Fig. 1E-H). Therefore, we assume that ZKO_C1 represents a hypomorph and ZKO_C2 a full knockout. Based on these validation experiments, LKO_C1 and LKO_C2 were designated as the mono- and biallelic *LMNA* knockout clones, respectively, and are hereafter referred to as LKO (0/+) resp. LKO (0/0). The ZKO_C1 and ZKO_C2 were designated as hypomorphic and biallelic *ZMPSTE24* knockout clones, respectively, and are hereafter referred to as ZKO (0/-), resp. ZKO (0/0).

### LMNA and ZMPSTE24 knockout differentially affect hiPSC-derived cardiomyocyte maturation

To evaluate the impact of the frameshift mutations on iCM development, we differentiated the hiPSC lines into iCM and cultured them as 2D monolayers up to 30 days post-differentiation (D30). As an initial readout of differentiation outcome, wells were categorized as non-beating, containing beating patches, or uniformly beating monolayers (Fig. 2A). We observed significant variability between the different iCM lines and between separate biological differentiations, likely due to the intrinsic heterogeneity of the iCM model (three biological replicates per cell line, 24-wells per differentiation). Unless stated otherwise, all subsequent experiments were restricted to cultures classified as beating monolayers. Monolayer density remained largely stable over time in CTRL and ZKO cultures. In contrast, LKO (+/-) and LKO (0/0) cultures showed a progressive reduction in cell density from D15 onward, with LKO (0/0) cultures displaying an almost complete loss of cells by D30 (Fig. 2B). We next assessed iCM maturation in the CRISPR-edited lines by comparing morphological and functional features. Cell size varied within and between phenotypes with ZKO (0/0) iCM showing significantly more compact shapes than CTRL cells. IF analysis further revealed reduced alpha-actinin levels and less organized sarcomere structures in both homozygous KO lines, suggesting less efficient structural maturation (Fig. 2C-E). To determine whether these structural abnormalities were accompanied by altered iCM function, we assessed intracellular calcium handling using Fluo-4 AM imaging (Fig. 2F). LKO (0/0) iCM exhibited increased beating frequency and higher calcium dynamic range, whereas ZKO (0/0) iCM showed significantly lower peak frequency and dynamic range compared with CTRL iCM (Fig. 2G-I). These findings indicate that loss of *LMNA* and *ZMPSTE24* both affect calcium handling and contractile behavior but do so in a distinct manner.

**Figure 2.**
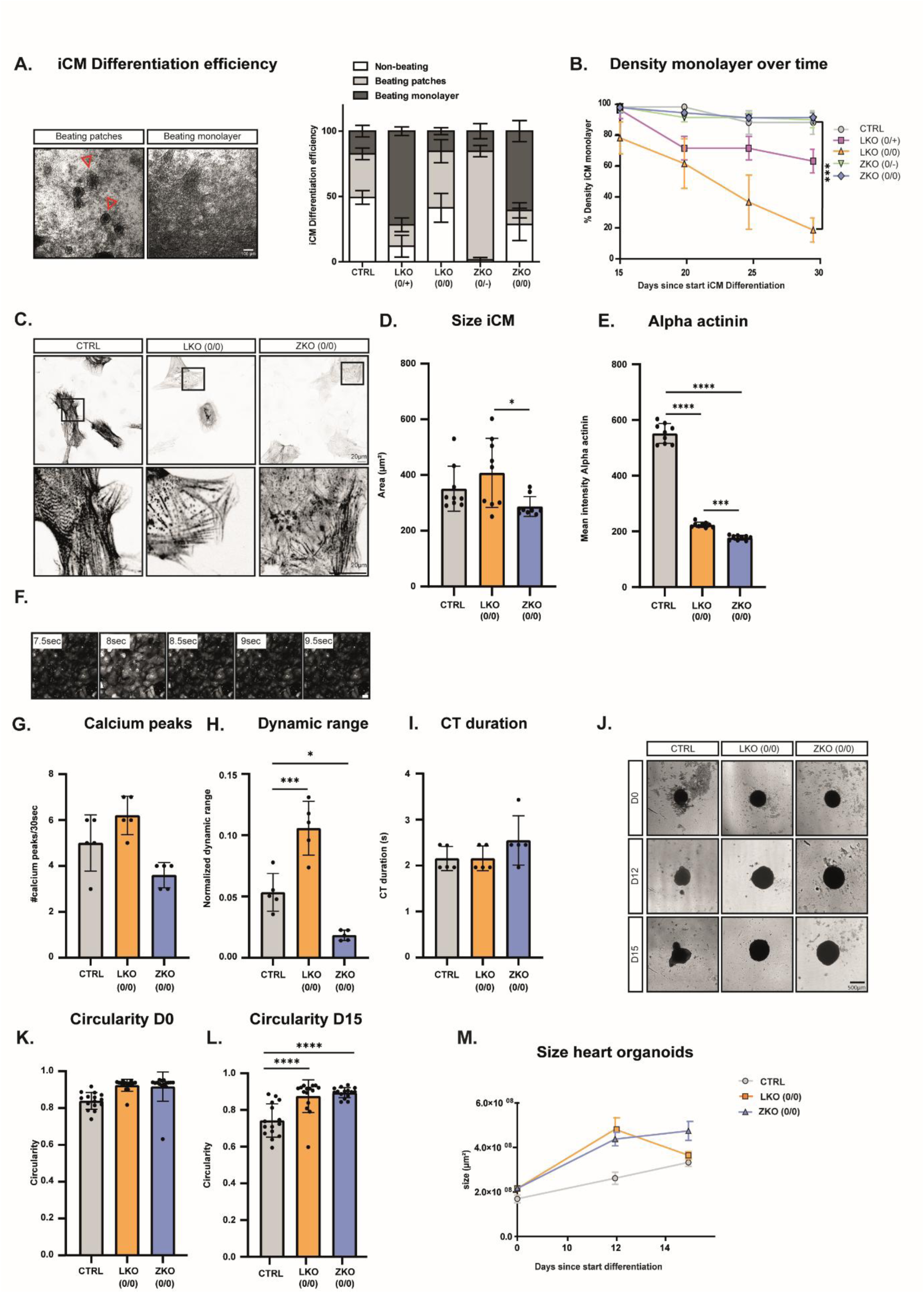
*LMNA* and *ZMPSTE24* knockout differentially affect hiPSC-derived cardiomyocyte maturation. (A) Brightfield image of iCM categorized as ‘beating patches’ and ‘beating monolayer”. Each well of 24well plate is categorized into one these three conditions, resulting in a percentage (n=3). (B) Quantification of the density monolayer over time on brightfield images (n_bio_=3, n_tech_=5) (C) Montage of representative images of immunofluorescence staining on D12 iCM for Alpha actinin. Zooms are contrast stretched. Scale bar = 20µm (D) Quantification of mean intensity immunofluorescence staining after segmentation cytoplasm using Cellblocks. (n_bio_=3, n_tech_=3, linear mixed model). (E) Quantification of cell area after segmentation cytoplasm using Cellblocks. (n_bio_=3, n_tech_=3, linear mixed model). (F) Montage of representative image of live cell imaging with FLUO4 AM staining. Scale bar = 50µM. (G) Detection and quantification of calcium peaks using in-house script (n_bio_=3, n_tech_=1-2, linear mixed model). (H) Quantification of normalized dynamic range of the calcium peaks. After bleach correction, the normalized dynamic range results from subtracting lowest point from highest point in calcium wave. (n_bio_=3, n_tech_=1-2, linear mixed model). (I) Quantification of average CT duraction. Measure time between detected begin and end of calcium wave. (n_bio_=3, n_tech_=1-2, linear mixed model). (J) Representative brightfield images of heart organoids on D0, D12 & D15 of differentiation. (K-L) Measurement circularity heart organoids on D0 and D15 of differentiation. (n_bio_=3, n_tech_=5). (M) Area of the heart organoids is followed up over time via brightfield imaging (µm^2^) (n_bio_=3, n_tech_=5).

To better model the three-dimensional physiological context, we generated self-organizing cardiac organoids (Fig. S2A-C). After optimization, we differentiated the bi-allelic knockout lines (three biological differentiations with at least 5 heart organoids per differentiation). All genotypes initially formed cardiac organoids of comparable size. However, LKO (0/0) cardioid organoids showed a marked reduction in size at D12, consistent with their density loss in 2D monolayer culture (Fig. 2J-M). In addition, CTRL organoids gradually lost their circular morphology and developed compartmentalized substructures, whereas both KO organoids retained a more circular shape throughout the culture period (Fig. 2K-M). Both LKO (0/0) and ZKO (0/0) organoids displayed reduced beating activity as compared to the CTRL (Fig. S2D&E). Together, these data show that *LMNA* and *ZMPSTE24* knockout differentially affect iCM differentiation state.

### LMNA and ZMPSTE24 knockout affect nuclear morphology in iCM

Given the central role of lamins in nuclear architecture, we next examined the impact of *LMNA* and *ZMPSTE24* KO on nuclear morphology during iCM differentiation (Fig. 3A). Visual inspection revealed a progressive deterioration of nuclear morphology. Classification of the nuclei into four distinct categories (‘ellipsoid’, ‘blebbed’, ‘elongated’, and allover ‘dysmorphic’) revealed that CTRL had the most normal, (‘ellipsoid’) nuclei, while loss of lamin A/C increased the percentage of ‘dysmorphic’ nuclei and accumulation of prelamin A comparatively increased the proportion of ‘blebbed’ and ‘elongated’ nuclei (Fig. 3C,D). Expansion microscopy revealed additional, subtler abnormalities including nuclear wrinkles and microblebs (Fig. 3B). To quantify these abnormalities over time, we measured nuclear circularity as a measure of roundness. At D-2, nuclear circularity was comparable across all cell lines but by D12, LKO (0/0) iCM showed a significant reduction in nuclear circularity compared with the other lines, consistent with their aberrant shape (Fig. 3F,G). Since circularity failed to capture the more subtle defects as observed in ZKO cells, we used a more sensitive contour-based analysis that extracts a curvature score from the sum of the first 50 elliptic Fourier descriptors (a series of harmonic coefficients describing the nucleus contour, where higher-order terms capture increasingly fine-scale deviations from a smooth, elliptical shape) (Fig. 3E) (21). This approach revealed significant differences in nuclear morphology across all cell lines at D12 and detected abnormalities in LKO (0/0) as early as at D2 (Fig. 3H,I). The delayed onset of nuclear abnormalities in the ZKO lines coincided with induction of (pre-)lamin A, which became detectable by IF from D8 onward (Fig. S1E). Together, these data indicate that *LMNA* and *ZMPSTE24* loss give rise to distinct nuclear phenotypes during iCM differentiation, with *LMNA* deficiency causing an earlier and more pronounced defect in nuclear shape, and *ZMPSTE24* deficiency leading to more gradual and subtle abnormalities.

**Figure 3.**
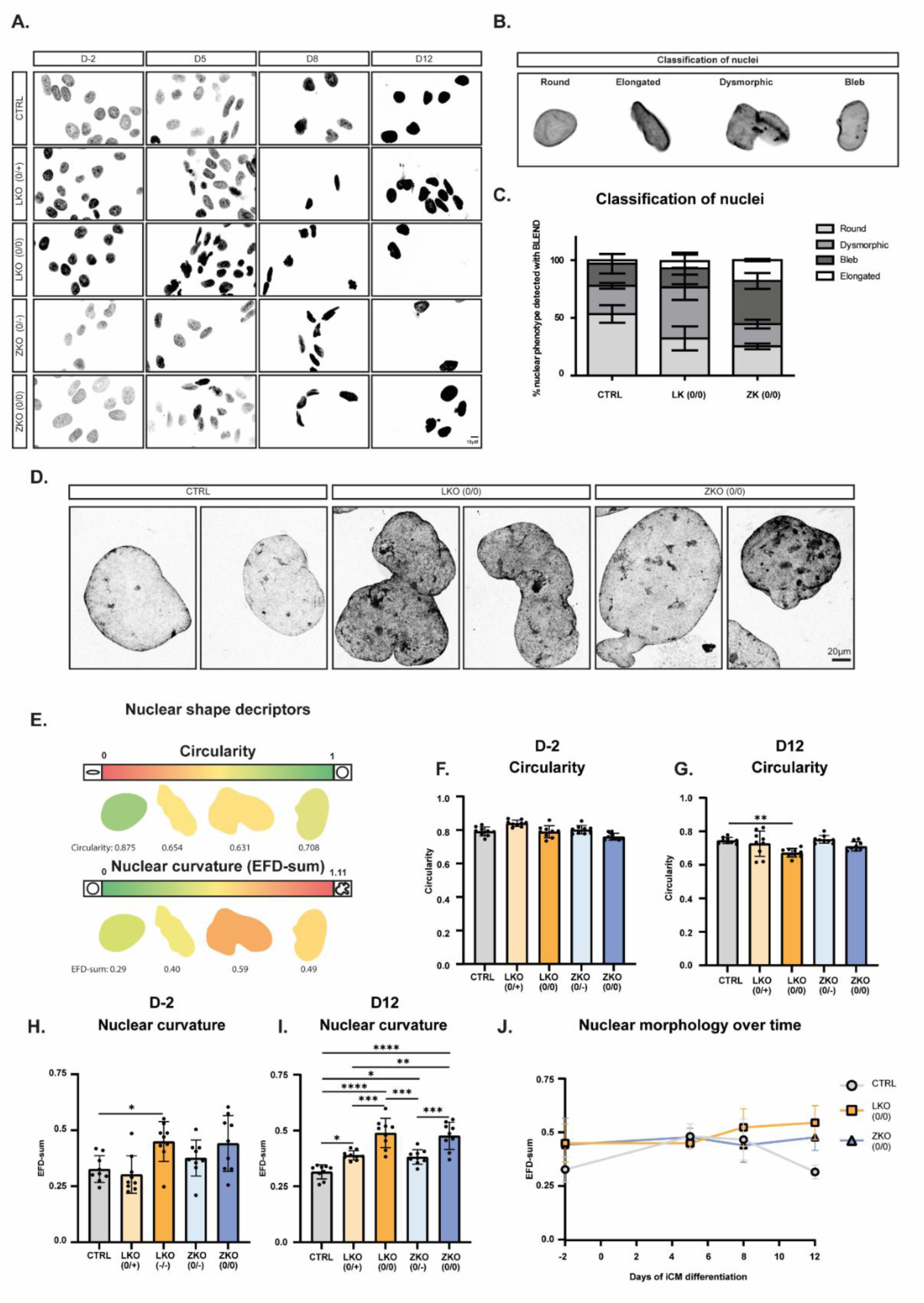
*LMNA* and *ZMPSTE24* knockout affect nuclear morphology in iCM. (A) Montage of representative images of DAPI staining on D-2, D5, D8 & D12 of iCM differentiation. Scale bar = 10µm. (B) D12 iCM magnified with expansion protocol and stained with Hoechst. Scale bar = 20µm. (C&D) BLEND (21) supervised classification of nuclei in four self-defined categories (round, elongated, dysmorphic, bleb). (n_bio_=3, n_tech_= 5 images). (E) Color coding of selected nuclei using color coder script for both circularity and nuclear curvature (EFD-sum) (47). (F&G) A dedicated script Cellblocks is used to detect nuclei in the nuclear DAPI channel. Circularity is quantified on D-2 and D12. (n_bio_=3, n_tech_=3 wells). (H&I) Quantification of the EFD-sum (sum of 50 EFD parameters) on D-2 and D12. (J) Follow nuclear morphology over time. EFD-sum for each timepoint (D-2, D5, D8, D12).

### Loss of LMNA and ZMPSTE24 induces distinct transcriptomic and proteomic remodeling

To identify molecular changes associated with complete loss of *LMNA* or *ZMPSTE24*, we performed bulk RNA sequencing (RNAseq) on LKO (0/0), ZKO (0/0) and CTRL iCM at D12. Dataset quality was supported by clustering of replicates from each genotype in a principal component biplot (Fig. S4A). Hierarchical clustering of the most dysregulated genes further confirmed clear segregation of the different lines (Fig. S4B). As expected, both CRISPR-targeted genes were strongly downregulated in the corresponding knockout lines (Fig. S4C). Using thresholds of log2-fold change ≥ 1, and Q value ≤ 0.05, we identified 1624 differentially expressed genes (DEGs) in LKO (0/0) vs CTR, 1755 in ZKO (0/0) vs. CTR and 2439 DEGs between ZKO (0/0) and LKO (0/0). Of these, 426 DEGs were shared between the two knockout lines (Fig. 4A), indicating both common and genotype-specific transcriptional responses. Interestingly, pathway enrichment analysis of the DEGs revealed a stronger representation of critical signaling pathways in LKO (0/0) such as HIF-1, Hippo and TGF-beta (with genes such as *HK1*, *PFKP* for the HIF-1 pathway and *CCN2*, *WWTR1* and *TGFB1* for the Hippo pathway and TGF-beta pathway respectively), whereas ZKO (0/0) showed a greater enrichment of pathways related to inflammatory signaling (Fc gamma R-mediated phagocytosis, leukocyte migration, inflammatory mediator regulation of TRP channels, Human papillomavirus infection pathway (typified by *e.g.* dysregulation of *TNF*, *CDC42* and *FN1s)*. Focusing on LKO (0/0)-specific DEGs, we identified several genes previously associated with DCM like *BAG3 and ACTA1* (22,23). Notably, *PPARα,* a central regulator of fatty acid metabolism and driver of lipotoxicity in DCM, was upregulated as well (24). Moreover, several of the top-ranked DEG, including *ADH1A* and *HMGCS2* further support the presence of a metabolic shift towards lipotoxic remodeling. *ACTN1, RHOA, MSN* are all connected to the actin cytoskeleton and were downregulated, suggesting structural cytoskeletal downregulation. Also, *KCNJ2*, encoding part of potassium channel stabilizing the rest membrane potential was found downregulated, and *MED25,* a mediator complex subunit involved in stress-response transcription, was found upregulated, consistent with recent reports connecting MED25 to Golgi stress in DCM (Fig. 4D). In contrast, ZKO (0/0)-specific DEGs included downregulation of *PLD5,* a phospholipase D family member involved in membrane-associated signaling and upregulation of *NPR1*, a receptor mediating natriuretic peptide signaling, suggesting differential engagement of stress-response and intracellular signaling in response to prelamin A accumulation. ZKO (0/0) cells also displayed upregulation of *HDAC5*, linking perturbed lamina formation to aberrant heterochromatin maintenance and transcriptional regulation (Fig. 4E). *ATM* and *DDX41* upregulation further indicate activation of DNA damage response and cytosolic DNA sensing pathways, linked to innate immune signaling cascades, whereas increased expression of *TRAF5* points to NF-κB-driven inflammatory signaling pathways. To complement the transcriptomic analysis, we performed proteomics on the same cell lines (Fig. S4B). Using thresholds of FDR < 0.2, and Log2-fold change of 1, we identified 1993 differentially abundant proteins (DAPs) in LKO (0/0) vs CTR, 1365 in ZKO (0/0) vs. CTRL, 668 DAPs were shared between the two knockout lines (Fig. 4F). Loss of lamin A/C resulted in broad proteomic remodeling across multiple cellular processes, including changes in RNA metabolism and translation, cellular organization and cytoskeletal dynamics (Fig. 4G & Fig. S4F). Interestingly, metabolic changes were also apparent here with the enrichment of carboxylic metabolic pathway and dysregulation of proteins involved in the fatty acid metabolism (e.g., *FABP1)*, strengthening the transcriptomic results (Fig. 4G & Fig. S4F). In agreement with the RNA-seq data, ZKO (0/0) cells showed stronger enrichment of immune- and inflammation-related proteomic signatures, the enrichment of multiple viral infection pathways describing a similar innate immunity inflammation pattern as seen in RNAseq (TBK1, DDX41, MAP3K7) (Fig. 4H & Fig. S4G). Together, these data demonstrate that *LMNA* and *ZMPSTE24* knockout induce distinct molecular remodeling programs in iCM, with *LMNA* loss more prominently associated with metabolic and structural pathway alterations, and *ZMPSTE24* loss linked to inflammatory and stress-related signaling.

**Figure 4.**
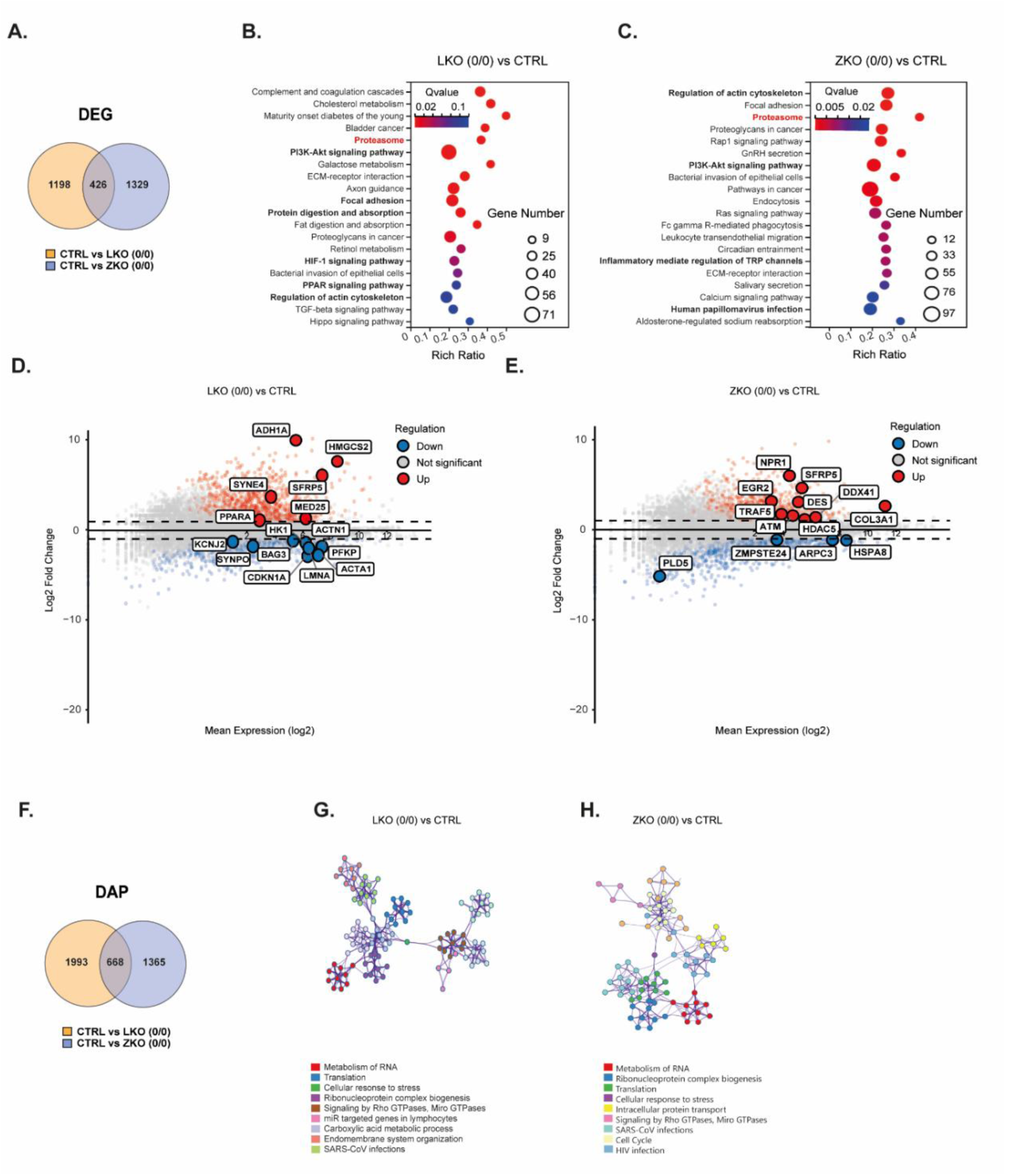
Loss of *LMNA* and *ZMPSTE24* induces distinct transcriptomic and proteomic remodeling. (A) Venn diagram show differential expressed genes (DEG) between CTRL vs LKO (0/0) and CTRL vs ZKO (0/0) with Qvalue < 0.05 and log2-fold change > 1. (B) Kegg enrichment bubble chart of DEGs LKO^−/−-^ vs CTRL. (C) Kegg enrichment bubble chart of DEGs LKO (0/0) vs CTRL. (D) MA plot showing interesting up- or downregulated genes (CTRL vs LKO (0/0)). (E) MA plot showing interesting up- or downregulated genes (CTRL vs ZKO (0/0)). (F) Venn diagram show differential abundant proteins (DAP) between CTRL vs LKO (0/0) and CTRL vs ZKO (0/0) with FDR < 0.02 and log2-fold change > 1. (G&H) Metascape network. Statistically enriched terms were hierarchically clustered into a tree based on Kappa-statistical similarities among their gene memberships (similar to what is used in NCI DAVID site). A subset of representative terms from the full cluster and converted them into a network layout. More specifically, each term is represented by a circle node, where its size is proportional to the number of input genes fall under that term, and its color represent its cluster identity (i.e., nodes of the same color belong to the same cluster). Terms with a similarity score > 0.3 are linked by an edge (the thickness of the edge represents the similarity score).

### Loss of LMNA and ZMPSTE24 converge on proteostatic defects

Several biological processes were shared between *LMNA* and *ZMPSTE24* knockout iCM, including PI3-AKT and focal adhesion pathways, suggesting a convergent disruption of cell survival signaling and cell-matrix interaction upon nuclear envelope perturbation. Notably, both KO models also exhibited a prominent downregulation of proteasome-related pathways. Functional annotation of DEGs and DAPs revealed a significant enrichment of pathways related to protein folding and protein degradation with several of the top enriched pathways shared between LKO (0/0) and ZKO (0/0) iCM (Fig. 5A-D). Among these, multiple proteasome components were coordinately downregulated in both knockout lines, suggesting an impairment of the ubiquitin-proteasome system system. (Fig. 5B&D). To assess whether these changes translated into compromised proteasome function, we measured 20S proteasome activity using a fluorigenic assay, with MG132-treated samples included as an inhibitory control (25,26). Consistent with the pathway analysis, proteasome activity was reduced in both knockout iCM lines, with the strongest reduction observed in ZKO (0/0) cells (Fig. 5E). We next examined whether the knockout iCM activated compensatory proteostasis pathways. Canonical autophagy-related genes, including ATG family members and LC3, were not upregulated at the transcript level and proteome level. Instead, we observed a conspicuous downregulation of *BAG3*, a key regulators of chaperone-assisted selective autophagy that has been implicated in DCM and in the shift from proteasomal degradation to autophagy under mechanical stress (Fig. 4D & Fig. 5F) (23,27). *BAG3* transcript levels were significantly reduced in LKO (0/0) (Fig. 5F), raising the possibility that BAG3 dysregulation contributes to the observed proteasome defects. This downregulation was confirmed at the protein level by quantitative IF, not only in LKO but also in ZKO. (Fig. 5F–H). Together, these data indicate that both lamin A/C loss and prelamin A accumulation impair proteasome activity in iCM and are not accompanied by clear induction of compensatory autophagy-related programs.

**Figure 5.**
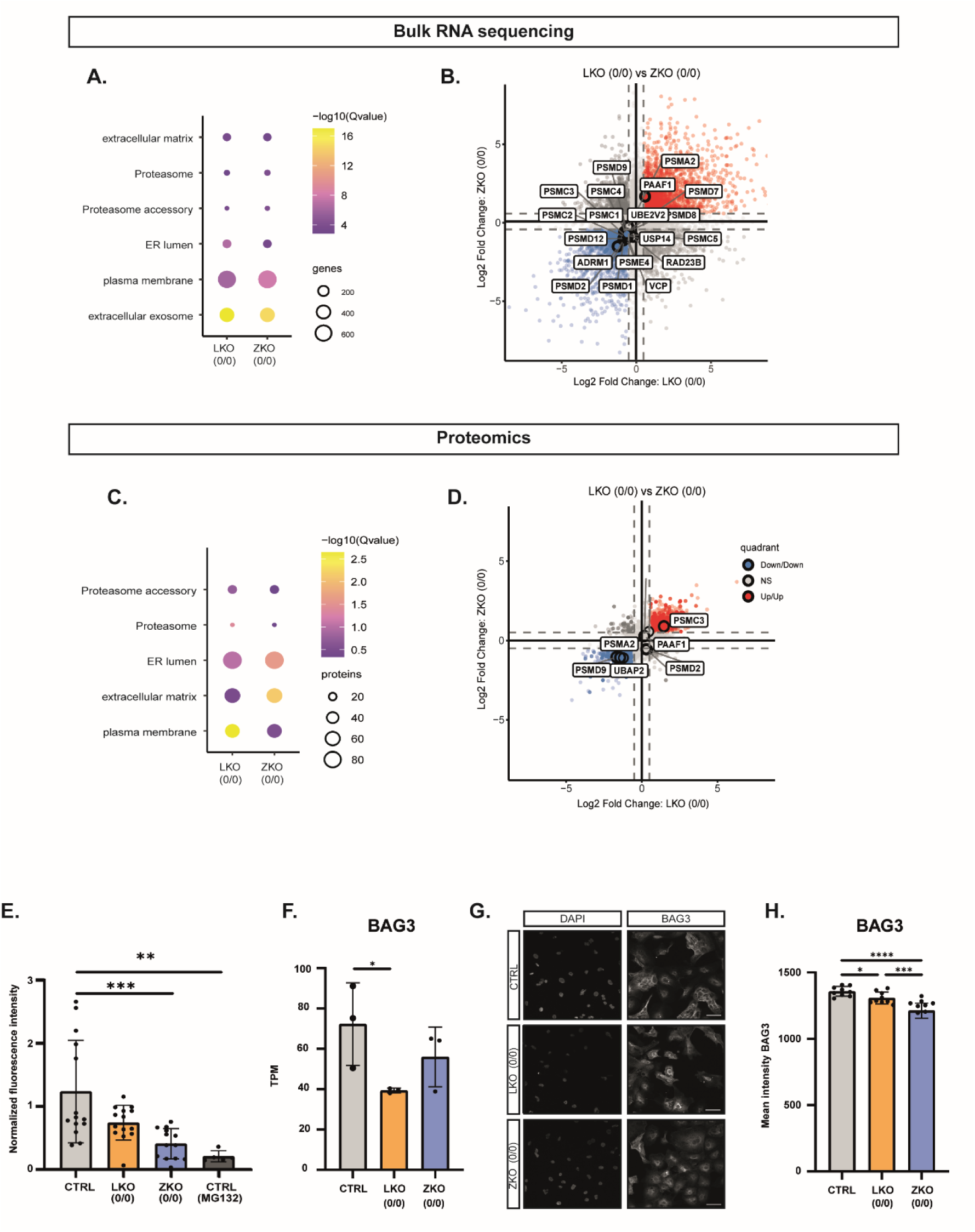
Loss of *LMNA* and *ZMPSTE24* converge on proteostatic defects. (A) Functional annotation DEGs based GO enrichment terms of bulk RNA sequencing. (B) Quadrant plot showing DEG (log 2 fold change) with CTRL vs LKO (0/0) on the X-axis and CTRL vs ZKO (0/0) on the y-axis. (C)) Functional annotation DAPs based GO enrichment terms of bulk proteomics. (D) Quadrant plot showing DAPs (log 2 fold change) with CTRL vs LKO (−/−-) on the X-axis and CTRL vs ZKO (0/0) on the y-axis. (E) Normalized measurement of proteasomal activity with 24h treatment with protease inhibitor MG132 as positive control. (n_bio_=3, n_tech_= 4-5wells, linear mixed model). (F) Quantification TPM for BAG3 (Qvalue < 0.05). (G) Montage of representative images of DAPI and BAG3 staining on D12 of iCM differentiation. Scale bar = 50µm. (H) Quantification of mean intensity immunofluorescence staining (BAG3) after segmentation cytoplasm using Cellblocks. (n_bio_=3, n_tech_=3, linear mixed model).

## Discussion

In this study, we generated and molecularly characterized iCM with partial or complete disruption of key determinants of A-type lamin metabolism, *LMNA* and *ZMPSTE24*. Although these perturbations converge on the nuclear lamina, they represent biologically distinct states: loss of lamin A/C removes an essential structural component of the lamina, whereas loss of *ZMPSTE24* leads to prelamin A accumulation, thereby altering lamin maturation and potentially introducing toxic gain-of-function effects. By comparing the models side by side, we show that disruption of lamin homeostasis in human iCM triggers both shared and divergent pathological responses, spanning nuclear architecture, calcium handling, metabolic and inflammatory signaling, and proteostasis.

A-type lamins are central to the maintenance of nuclear integrity in mechanically active tissues such as the heart. Previous studies have shown that disruption leads to nuclear deformation, impaired mechanical stability and susceptibility to NE damage under contractile stress (28–30). In line with this, LKO (0/0) iCM developed progressive abnormalities in nuclear morphology during differentiation, consistent with a loss of structural support as they acquire contractile function. The earlier onset and greater severity of the nuclear phenotype in *LMNA*-deficient cells further supports the notion that complete absence of lamin A/C destabilizes the nucleus more profoundly than defective lamin processing alone. Prelamin A accumulation in ZKO (0/0) was associated with subtler nuclear abnormalities that emerged later during differentiation, scaling with prelamin A accrual.

These structural differences were accompanied by clear functional divergence. LKO (0/0) iCM displayed increased CT amplitudes and a higher frequency of calcium peaks, pointing to dysregulated calcium cycling and electrophysiological instability. Such findings are in line with prior work linking *LMNA*-related DCM to abnormal calcium handling, altered excitability, and arrhythmogenic behavior and in line with our observed downregulation of *KCNJ2* (31,32). One plausible interpretation is that the remodeling of excitation-contraction coupling machinery contributes further to the disturbed mechanotransduction and disruption of the nuclear integrity, potentially through altered calcium handling and contractile dynamics impacting cytoskeletal tension. This may additionally promote actin cytoskeleton remodeling, characterized by reduced anchoring and retention at the membrane, thereby weakening mechanical coupling between the sarcolemma and intracellular cytoskeletal networks.

In contrast, ZKO (0/0) iCM showed reduced calcium dynamic range and lower beating frequency, suggesting a more hypocontractile or functionally immature state. This divergence indicates that defects in lamin metabolism cannot be viewed as a single pathological axis with a uniform downstream consequence. The progressive cell loss observed in LKO (0/0) cultures further suggests that, in the absence of lamin A/C, contractile stress on the nuclei may eventually exceed the capacity of the cells to maintain viability.

Our transcriptomic and proteomic analyses provide a broader molecular context for these phenotypes. Despite the limited overlap between the DEGs and DAPs, there was a strong concordance at the pathway level. In LKO (0/0) iCM, one of the most prominent signatures for transcriptomics and proteomics was the metabolic shift towards lipotoxicity *e.g.,* shown by the upregulation of *PPARα*, a major regulator of lipid uptake and mitochondrial β-oxidation (33). In the heart, PPARα signaling has also been linked to lipotoxic remodeling and contractile dysfunction (33,34). In the context of lamin A/C deficiency, this may reflect a maladaptive metabolic shift in response to sustained mechanical and nuclear stress. The concomitant induction of *MED25* further raises the possibility that Golgi-associated stress and altered secretory homeostasis contribute to this phenotype. Although these links remain correlative in the present study, they collectively support the view that *LMNA* loss has drastic consequences that extend beyond nuclear mechanics, reaching into core metabolic and organellar stress pathways.

By contrast, ZKO (0/0) iCM showcase a less pronounced phenotype, they were mainly characterized by several inflammation linked pathways showing the enrichment of the innate immune system, as exemplified by the upregulation of the DNA sensor *DDX41*, suggesting a activation of innate DNA sensing pathways. This is consistent with emerging evidence that defects in nuclear lamina integrity can promote cytosolic DNA sensing and type I interferon responses via cGAS-STING-dependent mechanism, as activation of this pathway may contribute to CM dysfunction and disease progression (35–37). Notably, in CM, lamin deficiency can lead to nuclear envelope rupture and DNA damage without triggering the cGAS-STING pathway due to insufficient accumulation of accessible cytosolic DNA (36).

Despite these differences, both models converged on focal adhesion, PI3K-AKT signaling pathway and most remarkable their shared defect in proteostasis, which may represent a central common denominator of lamin-associated cardiomyocyte dysfunction. Across transcriptomic, proteomic, and functional readouts, we observed coordinated downregulation of proteasome-related components and reduced proteasome activity in both *LMNA*- and *ZMPSTE24*-deficient iCM. This is a particularly relevant finding in CM, where long-lived proteins, high contractile demand, and limited regenerative potential place exceptional pressure on protein quality control systems (38,39). Reduced proteasome activity would be expected to impair basal turnover of damaged or misfolded proteins and increase vulnerability to cumulative stress during maturation and contraction. Within this shared proteostasis phenotype, BAG3 may represent an important mechanistic node. BAG3 is a critical regulator of cardiac protein quality control and a core component of chaperone-assisted selective autophagy (CASA), particularly in mechanically strained tissues (40). Other components of the CASA complex are also significantly reduced (*e.g., SYNPO* in LKO (0/0) and *HSPB8* in ZKO (0/0)) (Fig. 4D&E). The downregulation did not surface in our proteomics dataset, possibly due to the low abundance, near the detection limit. But the observation that *BAG3* transcript levels were significantly reduced in LKO (0/0) model, together with reduced BAG3 protein signal by quantitative IF, raises the possibility that BAG3 dysregulation contributes directly to the proteasome defects we observed. BAG3 has been implicated not only in autophagic clearance, but also in the broader integration of stress-responsive proteostasis pathways in the heart (41–43). Thus, rather than simply failing to engage a compensatory autophagic program, lamin-perturbed CM may be compromised at the level of a key regulatory hub that coordinates the transition between proteasomal degradation and alternative quality-control pathways. In that light, the combined suppression of proteasome-associated pathways and BAG3 may reflect a more profound collapse of adaptive proteostasis capacity, which could sensitize CM to both mechanical and metabolic injury.

Taken together, our findings support a model in which lamin homeostasis is required not only for nuclear architecture, but also for maintaining functional coupling between nuclear mechanics, intracellular signaling, and protein quality control in CM. Loss of lamin A/C appears to preferentially destabilize nuclear structure and promote calcium dysregulation and metabolic remodeling, whereas prelamin A accumulation more strongly engages inflammatory signaling. Yet, both perturbations converge on impaired proteostasis, highlighting this pathway as a potentially unifying vulnerability downstream of lamina dysfunction. This is consistent with broader evidence that dysfunction of the ubiquitin-proteasome system (UPS) is a common feature of cardiomyopathies, where it compromises protein quality control and promote disease progression (44,45). Notably, proteasome activity can be reduced in failing hearts without changes in the proteasome subunit levels, highlighting functional impairment rather than protein abundance (46). This convergence is particularly interesting from a therapeutic perspective, because it suggests that distinct upstream lamin defects may still share downstream liabilities that are more tractable to intervention.

This study has some limitations. First, although iCM are a powerful human model, they remain relatively immature compared with adult ventricular CM, which may influence the magnitude and nature of lamin-dependent phenotypes. Second, our analyses were largely performed in 2D monolayer cultures, which do not fully recapitulate the mechanical and architectural complexity of cardiac tissue. The cardiac organoids experiments already suggest that three-dimensional systems may capture additional aspects of disease progression, particularly with respect to tissue organization and survival, and they provide a valuable platform for future studies especially to repeat the bulk RNA sequencing and proteomic experiments. Third, the current data identify associations between lamina disruption, BAG3 dysregulation, and proteasome impairment, but do not yet establish direct causality. Follow-up experiments aimed at restoring BAG3 expression, modulating proteasome function, or interrogating nuclear stress signaling pathways such as cGAS-STING will be important for defining causal relationships more precisely.

In summary, this study establishes *LMNA* and *ZMPSTE24* knockout iCM as complementary human models of disturbed A-type lamin biology. By comparing complete lamin A/C loss with prelamin A accumulation, we show that these perturbations evoke distinct structural, functional, and molecular phenotypes while converging on a shared impairment of proteostasis. These models therefore provide a useful framework for dissecting how nuclear lamina dysfunction is translated into cardiomyocyte pathology and may help identify therapeutic strategies aimed at stabilizing proteostasis, preserving nuclear integrity, or interrupting maladaptive downstream signaling in lamin-associated cardiomyopathy.

## Resource availability

### Lead contact

Requests for further information and resources should be directed to and will be fulfilled by the lead contact, Winnok H. De Vos

### Materials availability

CRISPR/Cas9-edited hiPSC lines generated in this study will be made available upon request.

### Data and code availability

- Bulk RNA-seq and proteomics data have been deposited are publicly available as of the date of publication
- All original code is available at: https://github.com/DeVosLab
- Any additional information required to reanalyze the data reported in this paper is available from the lead contact upon request

## Acknowledgements

We thank Daniel Flender for helping with the analysis of the proteomics data. This research was performed with the support of Elien Theuns and Karen Sterckx.

## Author contributions

Conceptualization: WDV, MA, BL

Supervision & funding acquisition: WDV

Methodology: EGH, LO, MV, BV, LV

Experimental design & execution: LV, MV

Data analysis: LV, WDV

Software development: WDV

Writing: LV, WDV

## Methods

### Maintenance and validation of human iPSC lines

The hiPSC line used in this present study (iPSC0028, UNSPSC code: 12352200) was purchased from Sigma-Aldrich (Caucasian, female, 24 years old). IPSC0028 is hiPSC were cultured feeder free on Matrigel (BD Bioscience)-coated plates with Essential 8 medium (Thermo Fisher Scientific), maintained at 37°C in 5% CO_2_, and passaged weekly using ReLeSR (Stemcell technologies). During thawing, 10 µM Y-27632 (MedChem) was added for 24h. Mycoplasma was tested every two months (mycoplasma detection kit, VWR). To validate retention of pluripotency, we made use of an embryoid body (EB)-based differentiation assay. In brief, EB were formed from hiPSC in U-bottom 96-well plates. For 14 days, Essential 6 medium (Life Technologies) was replaced every two days to stimulate spontaneous differentiation of EB into the three germ layers. Thereafter, RNA was extracted, and RT-qPCR was performed with primers specific for each germ layer (Table 1).

**Table 1:**
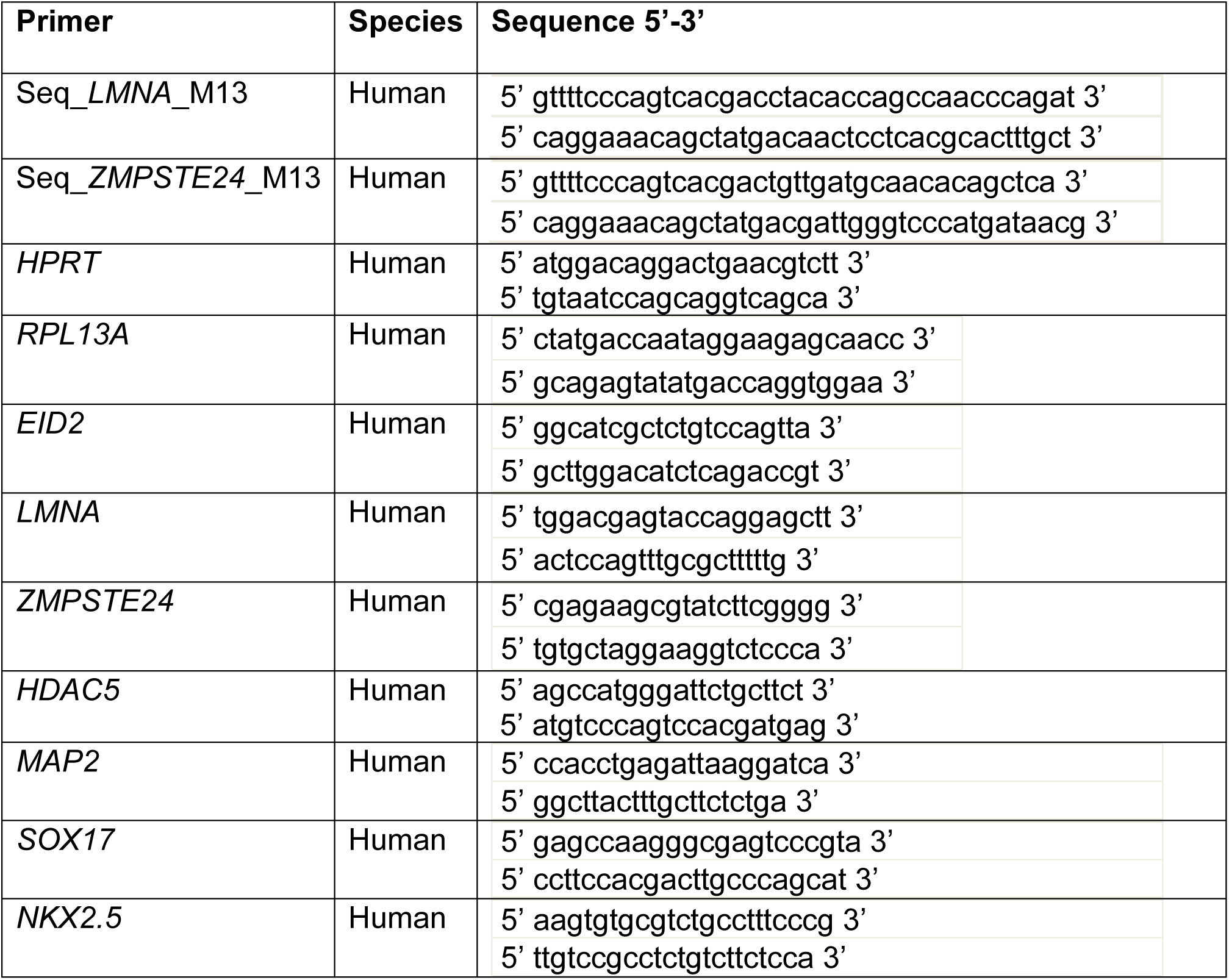
Overview primers sequences.

### CRISPR/Cas9 genome editing

Stable knockout iSPC0028 lines were produced using a CRISPR/Cas9 editing strategy based on a previously published protocol (48). The sgRNA sequences, targeting exon 1 of *LMNA* (5’ ccttcgcatcaccgagtctgaag 3’) and *ZMPSTE24 (*5’ ggccgagaagcgtatcttcgggg 3’) were designed with the CRISPR oligo design tool (Feng Zhang) (49). We selected sgRNAs with less than three predicted off-target site. Ribonucleoproteins (RNP) were prepared at a 7.5/1 sgRNA/cas9 (HiFi cas9 IDT) ratio and delivered to iPSC0028 (passage number 10) via nucleofection (program DS-138 of the 4D-nucleofector, Lonza). After nucleofection, hiPSC were left to recover in StemFlex medium (StemCell technologies) with 10 µM Y-27632 (MedChem express) and thereafter seeded at low-density. Colonies were picked, expanded clonally and DNA was extracted (QuickExtract, Biosearch Technologies). The target region of exon 1 of *LMNA* and *ZMPSTE24* was amplified by a Touchdown PCR using primers with M13-tag (FW_GTTTTCCCAGTCACGAC & RV_ CAGGAAACAGCTATGAC) adding a tail to simplify sequencing set-up. Sanger sequencing was performed using an ABI 3130XL Genetic Analyzer system (Applied Biosystems) according to standard protocol. Sequencing results were analyzed using ICE tool (Synthego) and DECODR, the latter using pair-end analysis.

### Cardiomyocyte Differentiation

Cardiac differentiation from hiPSC was adapted from a published protocol using small molecules to modulate the canonical Wnt signaling (50). To induce differentiation, hiPSC were seeded in 24-well plate (VWR) at 0.5×10^5^/cm^2^ density using TryplE (Life Technologies). The day of seeding is referred to as day minus two (D-2). After 48h, RPMI1640 medium (Thermo Fisher Scientific) with 2% B27 supplement without insulin (Life Technologies) and CHIR0092 (6µM, MedChem) was added for 24h to stimulate mesoderm formation (D0). On D3 of differentiation, RPMI1640 medium with 2% B27 supplement without insulin and iWP2 (5µM, SelleckChem) was added to the cells for 48h. Cells were subsequently cultured in RPMI1640 medium with 2% B27 without insulin until D7 of differentiation, on D7 switch was made to normal B27 supplement (Life Technologies) until D12 for validation experiments (referred to as iCM) and D30 for long culturing experiments. Biological replicates are separate differentiations. For RNA extraction and replating of iCM, dissociation was performed using TryplE 10x (Life Technologies) according to the supplier’s protocol with RPMI1640 medium supplemented with 1/1000 Y-27632 (MedChem express), 2% B27 supplement, 10% knock-out serum (Life technologies).

### Heart organoid differentiation

To differentiate hiPSC into heart organoids, we used the same differentiation protocol as described above on EB with some adaptations. hiPSC were seeded at a density of 10.000/well in U-bottom 96-well plates (VWR) and centrifuged at 100g, 3 min. Differentiation started 48h later, with 24h incubation of 6µM CHIR99021 in RPMI1640 medium with BMP4 (1µM) and Activin A (1.5ng/ml). On D2 of the differentiation, Wnt-C95 (5µM, MedChem) and L-ascorbic acid (200µM) were added for 48h. On D7, an extra incubation of 1h incubation CHIR0092 was performed. The heart organoids were imaged on fixed timepoints to follow the growth on an inverted brightfield microscope.

### Immunofluorescence staining

Cells were fixed with 4% PFA for 25min followed by 3x PBS wash step (Life Technologies, 14190-169). Thereafter, blocking solution phosphate buffered saline (PBS with 0.05% Thimerosal (Merck), 0.01% NaN_3_ (Merck), 0.3% bovine serum albumin (Thermo Fisher Scientific), 10% H1270 (Thermo Fisher Scientific)) with 10% Triton-X 100 (Sigma) was added for 45 minutes. Primary antibodies (Table 2) were prepared in blocking solution and added overnight at 4°C. After 3×5 min wash step with PBS, the secondary antibodies (Table 2) were added for 2h. Afterwards, cells were washed again with 3×5 min PBS and stained with DAPI (1µg/ml) for 15 min.

**Table 2:** Overview antibodies.

| Ab | Species | Vendor | Catalog number | dilution |
| --- | --- | --- | --- | --- |
| **Primary antibodies** |  |  |  |  |
| Prelamin A | Goat | Santa Cruz<br>Biotechnology | Sc-6214 | 1:200 |
| Lamin A/C | Mouse | Santa Cruz<br>Biotechnology | sc-376248 | 1:300<br>(WB<br>1:1000) |
| Lamin B1 | Rabbit | Abcam | ab16048 | 1:300 |
| Alpha-actinin | Mouse | Sigma | A7811 | 1:300<br>(WB<br>1:1000) |
| Cardiac Troponin I | Rabbit | Abcam | Ab47003 | 1:300 |
| Ubiquitin | Mouse | Cell Signaling<br>Technology | 3936 | 1:200 |
| BAG3 | Rabbit | Proteintech | 10599-1-AP | 1:150 |
| **Secondary Antibodies** |  |  |  |  |
| Cy<sup>TM</sup>3 AffiniPure®<br>Anti-Rabbit IgG | Donkey | Jackson<br>ImmunoResearch | 711-165-152 | 1:1000 |
| Alexa Fluor® 488<br>AffiniPure® Anti-Mouse IgG | Donkey | Jackson<br>ImmunoResearch | 715-545-150 | 1:1000 |
| Cy™5 AffiniPure®<br>Anti-Goat IgG | Donkey | Jackson<br>ImmunoResearch | 705-175-147 | 1:1000 |
| Anti-Mouse IgG<br>Polyclonal<br>Antibody (CF™<br>488A) | Donkey | Biotium | #20014 | 1:200 |
| Anti-Rabbit IgG<br>Polyclonal<br>Antibody (CF™<br>568) | Donkey | Biotium | #20098 | 1:200 |
| Anti-Goat IgG<br>Polyclonal<br>Antibody (CF™<br>633) | Donkey | Biotium | #20127 | 1:200 |
| HRP-conjugated<br>anti-mouse | Goat | Sigma-Aldrich | A4416 | 1:5000 |
| HRP-conjugated<br>anti-rabbit | Goat | Sigma-Aldrich | A6154 | 1:5000 |

Heart organoids were fixed on D15 of differentiation (D15) overnight in 4% paraformaldehyde (PFA, Carl Roth) and used for further experiments. Thereafter, they were washed with PBS and separately embedded in 2-4% agarose (Invitrogen) and sliced into 80µm thick slices using a vibratome. Next, the vibratome slices were dried on SuperFrost slides on 37°C for 1h.

### Westen blot

On D12 of differentiation, iCM were lysed using ice-cold RIPA buffer (freshly made, 2% SDS). Protein concentration was measured with Pierce^TM^ BCA Protein Assay kit (Life Technologies). Lysates (70%) were then mixed with NuPAGE LDS sample buffer (25%, Life Technologies) and dithiothreitol (DTT, 5%, Life Technologies) and heated for 5min at 95°C. Thereafter samples were loaded on the NuPAGE Novex 4-12% Bis-Tris Protein Gels (Life Technologies) with MOPS running buffer (Life Technologies) for 1h30 at 130V. Next, proteins were transferred to PVDF membranes (0.2µm pore size, Life Technologies) using a transfer mix of NuPAGE Transfer buffer (Life Technologies), methanol and NuPAGE transfer buffer (Life Technologies) for 1h, at 30V. Thereafter, membranes were blocked using blocking buffer (5% ECL in Tris Buffered Saline with 0.2% Tween 20 (TBST)) and subsequently incubated overnight (ON) at 4°C with primary antibodies, diluted in blocking buffer (Table 2). Horse radish peroxidase (HRP)-conjugated goat anti-mouse (Sigma-Aldrich A4416) and HRP-conjugated goat anti-rabbit (Sigma-Aldrich) were used as secondary antibodies. Chemiluminescence was used to detect the proteins on the membranes (Clarity^TM^ Western ECL Substrate) using a western blot imager (Cytiva, ImageQuant800). Quantification was done using the ImageQuant software (Cytiva) with GAPDH and Amersham^TM^ QuickStain Protein Labeling kit (Cytiva) for reference.

### qPCR

RNA was extracted from iCM on D12 of differentiation using miRNeasy kit (Qiagen), followed by cDNA synthesis using iScript^TM^ cDNA synthesis kit (Bio-rad). Thereafter RT-PCR was performed (40 cycles of 95°C for 30 sec, 95°C for 15 sec and 60°C for 30 sec and a melt curve ranging from 65 – 95°C with 0.5°C increment every step of 3 sec) using primers listed in Table 1. Reference genes were tested and the most stable 3 were selected (*HPRT*, *EID2*, *RPL13A*).

### RNA Sequencing

The iCM samples (three biological replicates for all five KO hiPSC lines) were harvested at the appropriate timepoint (D12) and RNA was extracted using the miRNeasy kit (Qiagen). RNA concentration and integrity (RIN) were assessed using a Bioanalyzer (Agilent). Additional RNA quality control was performed by BGI genomics, samples with total RNA >200ng, RIN > 7 and 28S/18S>1 were included for library preparation and further sequencing at BGI genomics (RNAseq, oligo dT, stranded, DNBseq platform, PE150, 20M reads per sample). A total of 15 iCM samples were tested and analyzed using the DNBseq platform. The average alignment ratio of the sample comparison (Homo_sapiens_NCBI_GCF_000001405.40_GRCh38.p14) was 98.58%, which is considered high. The average alignment of the gene set was 82.28%; a total of 18836 genes were detected. Quality filtering was performed with the SOAPnuke software v2.3 (BGI genomics), > 80% of raw reads were retained as clean reads across samples. Clean reads exhibited high base quality with Q20 values exceeding 98% and Q30 values above 92%, yielding approximately 6.62G data per sample. HISAT2 v2.0.4 and Bowtie2 were used to align clean reads to the reference genome and genes (https://www.ncbi.nlm.nih.gov/datasets/genome/GCF_000001405.40/) resulting in a uniquely mapped read alignment of >88%. Thereafter, gene quantification analysis and other analysis based on gene expression (principal component analysis, correlation and differential gene screening) were performed using R package-function princomp and DEseq2. DEG were defined as genes with log2-fold change ≥ 1, and Qvalue ≤ 0.05. All analysis and the generation of figures were performed in the Dr. Tom software (BGI genomics).

### Proteomics

iCM samples were harvested at the appropriate timepoint (D12) and pellets were collected (3 biological replicates for CTRL, LKO (0/0) and ZKO (0/0) iCM). Protein samples were further processed using micro S-Trap columns according to the manufacturer’s protocol, with the inclusion of iodoacetamide for alkylation (Profiti). Samples were digested with trypsin, and the resulting peptides were loaded onto Evotips according to the manufacturer’s instructions (Evosep). Peptide separation prior to mass spectrometry analysis was performed using an Evosep One system equipped with an EV1137 column, 15 cm × 150 µm, 1.5 µm particle size (Evosep). The column was coupled online to a timsTOF Pro mass spectrometer operating in positive ion mode and equipped with a CaptiveSpray ion source (Bruker Daltonik GmbH, Bremen, Germany). The timsTOF Pro was calibrated according to the manufacturer’s guidelines. The ion transfer capillary temperature was set to 180°C. Data were acquired using a Parallel Accumulation–Serial Fragmentation data-independent acquisition method, dia-PASEF ((51). The dia-PASEF window scheme covered an m/z range of 400–1200 and a 1/K0 range of 0.6–1.6, using 32 × 16 Th windows with a ramp time of 100 ms. The acquired raw files were exported and processed using PEAKS Online 13. Relative quantification was performed using label-free quantification. Database searching was performed using a target-decoy strategy against the human UniProt database, with the false discovery rate set to 1%. Trypsin was specified as the digestion enzyme, allowing up to two missed cleavages. Carbamidomethylation was set as a fixed modification, whereas deamidation and oxidation were set as variable modifications. Statistical analysis was performed using BigOmics Playground. Protein intensity values from PEAKs Online were normalized using the max-median normalization method. Missing values were imputed using singular value decomposition (SVD)-based imputation. Differential abundance analysis was performed between CTRL and experimental groups, and results were visualized with MA plots and Metascape.

### Expansion microscopy

Expansion microscopy was performed according to the Magnify protocol (52), with adaptations to optimize expansion for cells on coverslips. 12 mm glass coverslips (VWR, 631-1577) were placed in a 12-well plate, sterilized by spraying with 70% ethanol, air-dried, and washed twice with sterile PBS (Life Technologies, 14190169). Cells were then seeded onto the sterilized coverslips at a density of 90,000 cells per well and allowed to adhere overnight. Cells were then treated according to the experimental design, fixed in 2% PFA (Roti-histofix 4% paraformaldehyde, Roth, 3105.2) for 20 min, and stained using previously described immunolabeling procedures. After staining, samples were stored in PBS containing sodium azide until further processing. The monomer solution, freshly prepared for each experiment, consisted of 4% DMAA (Sigma-Aldrich, 274135-100ML), 34% sodium acrylate (Santa Cruz Biotechnologies, sc-236893), 10% acrylamide (Bio-Rad, 1610140), 0.01% N,N′-methylenebisacrylamide (Sigma-Aldrich, M7279-25G), 1% NaCl (Merck, S9625-1KG), and 1× PBS. Immediately before embedding, the monomer mix was supplemented with 0.25% ammonium persulfate (Sigma-Aldrich, A3678-25G), 0.05% TEMED (Sigma-Aldrich, T9281-25ML), and 0.1% methacrolein (Sigma-Aldrich, 133035-25ML) and mixed thoroughly. Gelation was carried out in custom-made chambers by placing silicone isolator sheets (Sigma Aldrich, GBL664581-5EA) with self-punched holes onto glass microscope slides. Stained coverslips were positioned cell-side down in direct contact with the monomer solution, and polymerization was allowed to proceed overnight at 37°C in a humidified environment. Following gelation, coverslips were removed and gels transferred to 6-well plates for denaturation. Homogenization was carried out in buffer containing 20% SDS (Sigma-Aldrich, L3771-1KG), 8 M urea (Sigma-Aldrich, U5378-500G), 25 mM EDTA (Thermo Fisher Scientific 15575020), and 2× PBS. Samples were incubated for 6h at 80°C in a humidified chamber, after which gels were washed three times in 1× PBS at room temperature and subsequently incubated for 1h in 1% C12E10 (Sigma-Aldrich, P9769-500G) at 60°C to remove residual SDS. Gels were then washed three additional times in PBS for at least 10 min each and stored overnight in PBS at room temperature before expansion. For expansion, gels were transferred to excess deionized water and allowed to expand, with repeated water exchanges until maximal expansion was reached. Expanded samples were stained with Hoechst (Invitrogen™, H3570, 1:1000) for 30 min, washed thoroughly with water, and imaged immediately to ensure optimal gel stability and fluorescence retention. Imaging was performed in poly-L-lysine (Sigma-Aldrich, P8920-100ML) coated glass-bottom dishes (CELLviewTM, Greiner Bio-One, 627860) to minimize gel drift. Expansion isotropy and expansion factor were validated using Gelmap (53).

### Proteasome activity measurement

Proteasomal activity was measured using the Proteasome 20S activity assay kit (Abcam, ab112154). iCM were seeded on D12 on Matrigel (BD Bioscience) coated on Nunclon^TM^ Delta Surface plates (Thermo Fisher Scientific) with a density of 8000 iCM/well. After 24h, medium was replaced and control compounds were added to selected wells. The proteasome inhibitor MG132 (Tocris Bioscience, 1748) was used as a positive control at a concentration of 2µM for 24h (54). At 47h, 100µl proteasome assay solution was added supplemented with Hoechst (20ng/ml) to normalize for cell density and incubated for 1h at 37°C and 5% CO_2_. Fluorescence was measured using the VANTAstar Microplate reader (BMG Labtech) with excitation/emission set at 490/520nm for R110 and 340/480nm for Hoechst.

### Microscopy

Live cell calcium imaging was performed on a fully automated Nikon CSU-W1-01 SoRa spinning disk confocal microscope, mounted on a Nikon Ti Eclipse body equipped with Perfect Focus System and a microscope incubator equilibrated at 37°C and 5% CO_2_. iCM were seeded in 96-well plates (Greiner) 48h before imaging and stained with Fluo-4 AM (Life Technologies) and Spy-DNA (Spirochrome) for 30 min with a subsequent 30 min waiting step. Recordings were made every 500ms for 30sec for each selected region using 20x/0.75 Plan Apo Dry lens. 488nm laser (excitation of Fluo-4 AM) and 640 nm (excitation of Spy-DNA) diode lasers were used in conjunction with a quad-band beam splitter and 520/35 nm and 685/40 nm bandpass emission filters, respectively. Images were acquired with a Prime 95B camera (Teledyne Vision Solutions), 1 central point/well with 5 wells per condition.

The same microscope was used for confocal recordings of immunostained cells (hiPSC and iCM) and heart organoid sections. At least three wells were used as technical replicates for each condition. Per well, 26 regions were imaged, using 40x/0.95 Plan Fluor Dry lens or 20x/0.75 Plan Apo Dry lens. 405 nm (DAPI), 488 nm (FITC), 560 nm (Cy-3) and 640 nm (Cy-5) diode lasers and 447/60 nm, 520/35 nm, and 685/40 nm bandpass filters were used. Images were acquired using a Kinetix sCMOS camera (Teledyne Vision Solutions). For vibratome sections, 1 image was taken per slice of heart organoid with 20x/0.75 Plan Apo Dry lens.

To quantify the survival of the iCM over time, we used the widefield inverted Nikon Ti microscope in brightfield mode. Three points of each well were selected using brightfield with 4x/0.1 Dry lens. On D12, the beating frequency was measured at a rate of 20 frames per second.

To follow the growth of the heart organoids over time, we used the widefield inverted Nikon Ti microscope in brightfield mode. The center of each well was automatically imaged at fixed timepoints using brightfield with 4x/0.1 Dry lens. On D15 of organoid differentiation or 2D iCM differentiation, images of contracting cells were acquired at a rate of 20 frames per second.

ExM images were acquired on a spinning-disk confocal microscope (Nikon CSU-W1 SoRa) with a 40× 1.2 NA objective (Apo 40x WI λS DIC N2) and a Kinetix sCMOS camera at a resolution of 0.16 µm/pixel.

### Image Analysis

Fluorescent image analysis was performed in FIJI image analysis freeware. A dedicated script Cellblocks (55) ((https://github.com/DeVosLab/CellBlocks) was used to detect nuclei in fixed assays and live cell imaging in the nuclear counterstain channel (DAPI or Spy-DNA) using a trained convolutional neural network as implemented in the StarDist plugin. Both morphological and intensity measurements were extracted from detected nuclear and cytoplasm regions. To measure the nuclear morphological and textural parameters BLEND was used (21). Elliptic Fourier Descriptors (EFD-sum) and entropy were used as parameters.

For live cell calcium imaging, as intensity fluctuations were measured across the full field of view and a dedicated script was used to extract relevant activity metrics (CalciumAnalysis.ijm on Github/DeVosLab).

### Statistics

Statistical analysis was conducted in R (56). To compare conditions, linear mixed models with independent biological replicate as random factor and technical replicate (mostly wells for quantification immunofluorescent staining) as nested factor were used. This was combined with Tukey *post hoc* method to control for errors during multiple comparisons. Adjusted p-values < 0.05 were considered significant. Graphs were generated with Graphpad Prism 10 and Rstudio. Details on the number of biological replicates, statistical methods and results are mentioned in the figure legends.

