## Supplemental Figure S1 for "CRISPR-Engineered hiPSC-derived Cardiomyocytes Reveal Divergent Responses to Loss and Defective Processing of A-type Lamins"

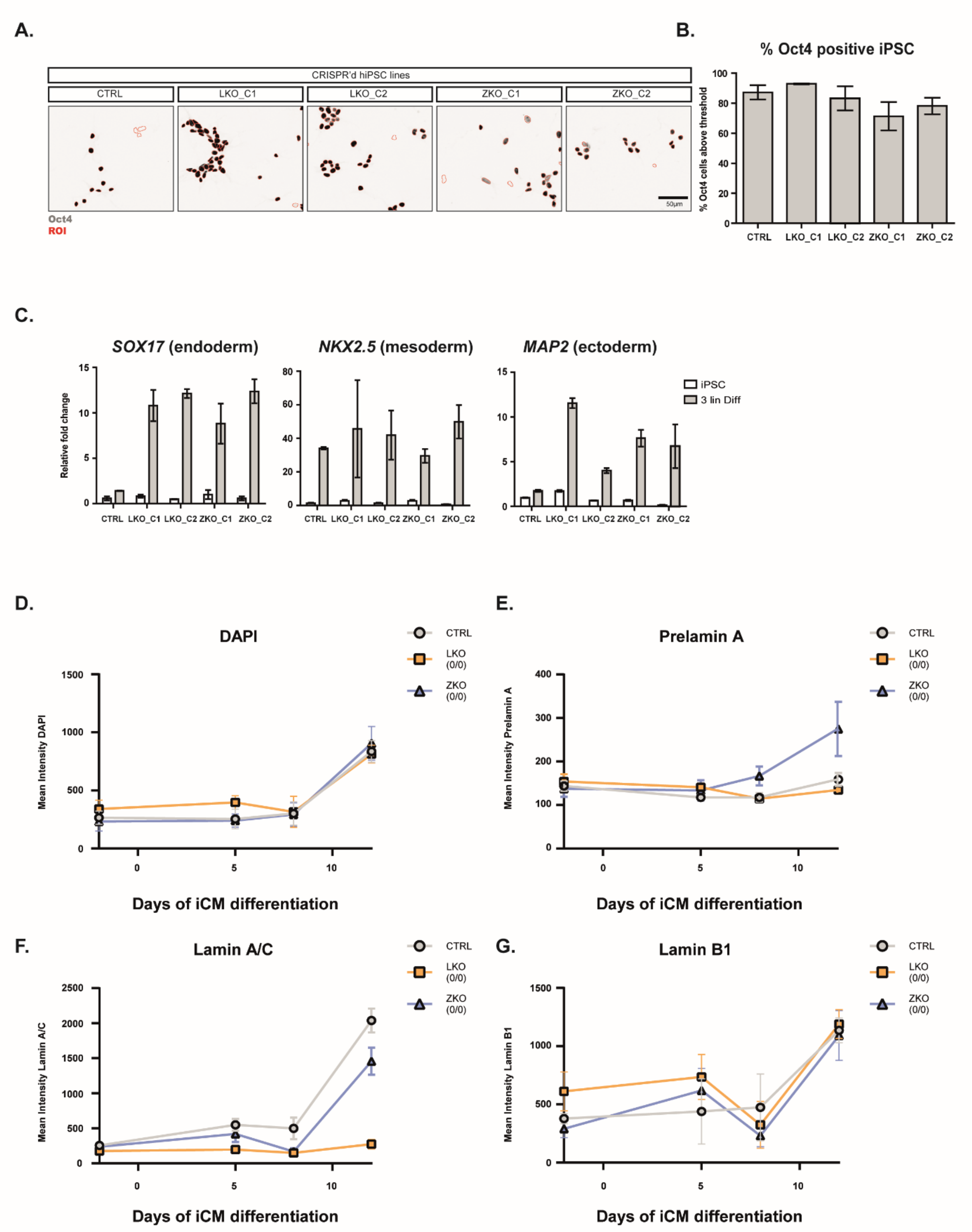


**Figure S1. Validation pluripotency CRISPR-edited hiPSC lines.**

(A) Montage of representative images of immunofluorescence staining for DAPI and Oct4. A dedicated script Cellblocks is used to detect nuclei in the nuclear DAPI channel. Detected nuclei are represented as region of interest (ROI). (B) Quantification of % Oct4 positive cells (threshold = 180, n=3) before iCM differentiation. (C) Three lineage differentiation is performed on CRISPR'd hiPSC lines. Quantification of relative fold change from qPCR analysis performed with primers *SOX17*, *NKX2.5* & *MAP2* (marker for endo-, meso- and ectoderm) (n=3). (D-G) Quantification of mean intensity immunofluorescence staining (DAPI, prelamin A, lamin A/C & lamin B1) after segmentation nucleus using Cellblocks at timepoint D-2, D5, D8 & D12 of iCM differentiation. ($n_{bio}$=3, $n_{tech}$=3).
