## Supplemental Figure S2 for "CRISPR-Engineered hiPSC-derived Cardiomyocytes Reveal Divergent Responses to Loss and Defective Processing of A-type Lamins"

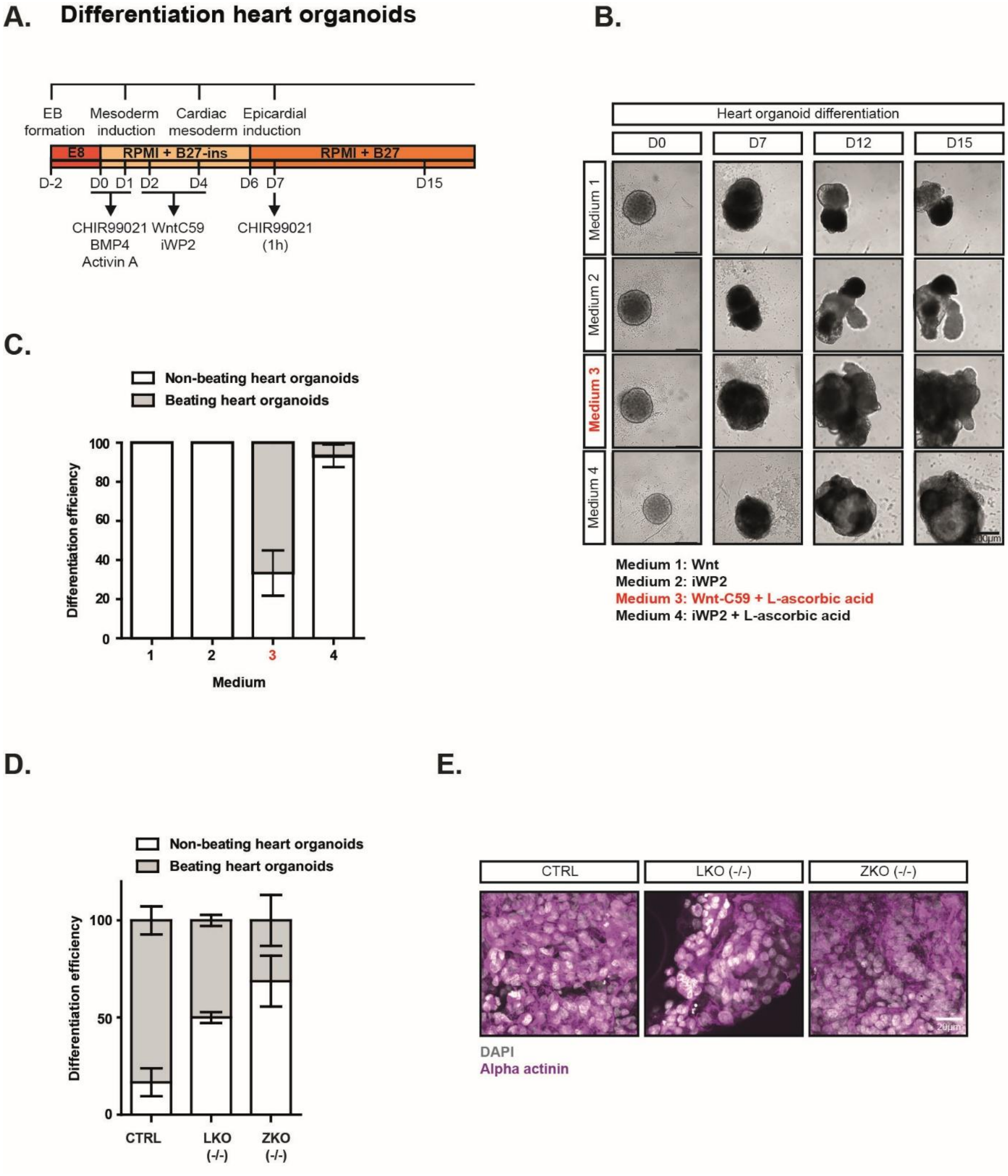


**Figure S2. 3D differentiation of CRISPR-edited hiPSC lines**

(A) Schematic overview heart organoid differentiation. (B&C) Optimization heart organoid differentiation protocol. Representative brightfield images of heart organoids. Four different differentiation protocols are compared (represented as medium 1- 4). Efficiency of differentiation is determined on D12 of differentiation. ‘Beating heart organoid’ = contraction seen during 1 min imaging. ‘Non-beating heart organoid’ = no contraction seen during 1 min imaging ($n_{bio}$=3, $n_{tech}$=5). (D) Efficiency of differentiation (‘beating’ or ‘non-beating’) is determined on D12 of differentiation ($n_{bio}$=3, $n_{tech}$=5). (E) Vibratome coupes are stained for DAPI and alpha actinin.
