## Supplemental Figure S3 for "CRISPR-Engineered hiPSC-derived Cardiomyocytes Reveal Divergent Responses to Loss and Defective Processing of A-type Lamins"

A.

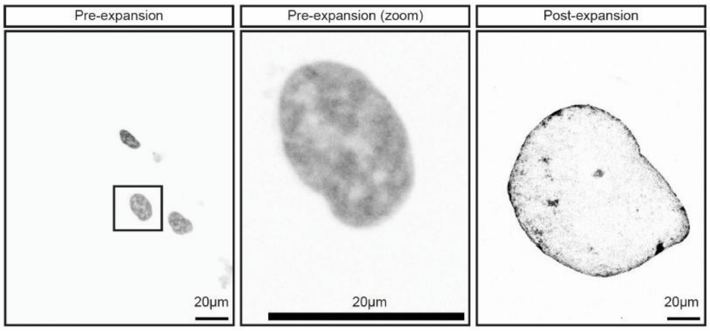


**Figure S3. Expansion protocol CRISPR’d iCM.**
**(**A) Montage of representative individual CTRL iCM . Left panel = DAPI staining on D12 CTRL iCM. Middle panel = Zoom in on individual nucleus from left panel. Right panel. Post-expansion image from individual CTRL iCM nucleus stained with Hoechst.  Scale bar=20µm.
