## Supplemental Figure S4 for "CRISPR-Engineered hiPSC-derived Cardiomyocytes Reveal Divergent Responses to Loss and Defective Processing of A-type Lamins"

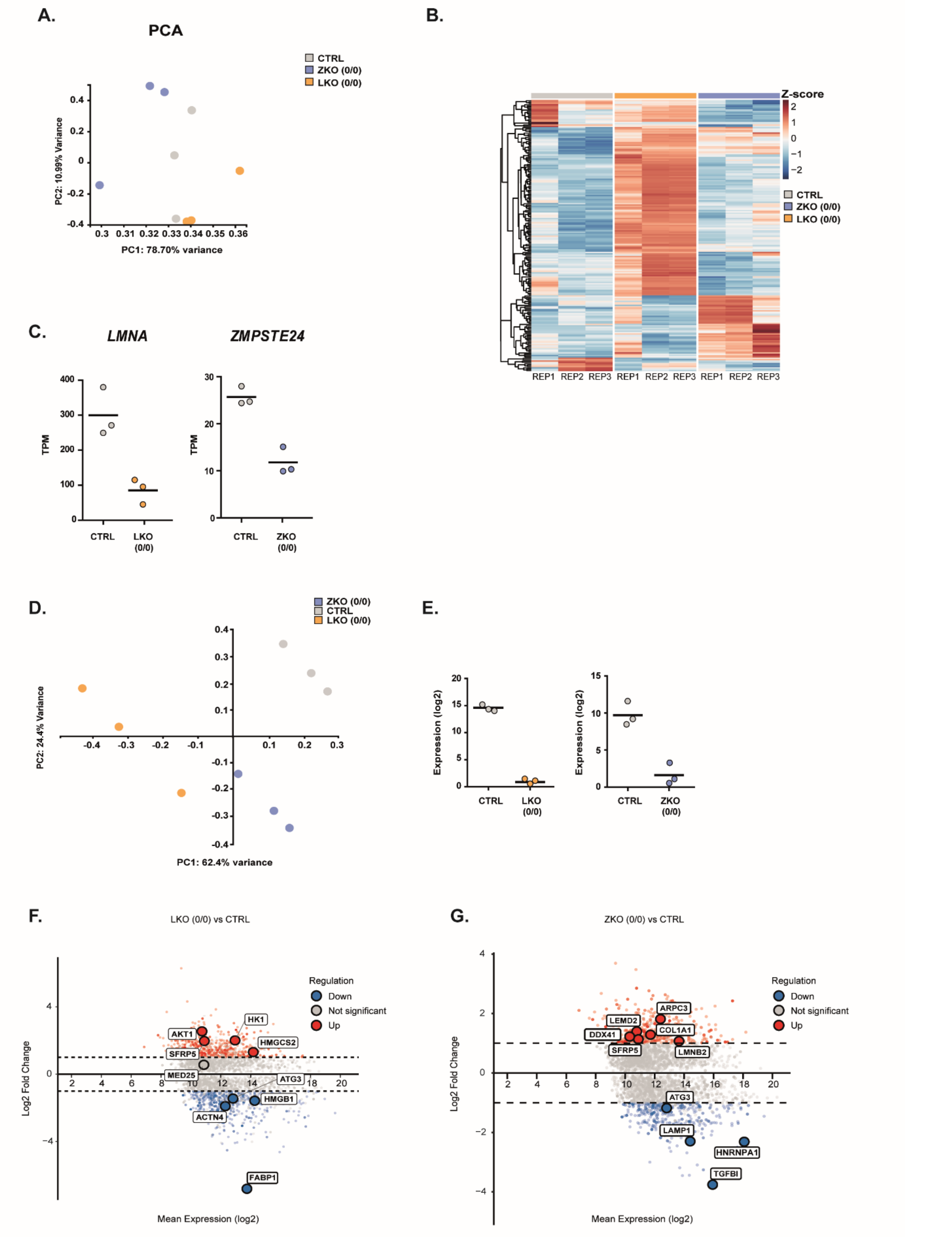


**Figure S4. Bulk RNA sequencing & proteomics.**
(A) Clustering of individual replicates of bulk RNA sequencing per line after principal component analysis (PCA) of all the read counts according to the first principal component, which accounted for 78.70%. (B) Heatmap of the top variable differentially expressed genes (DEGs) across CTRL, LKO (0/0), and ZKO (0/0), showing log2-transformed, row-scaled expression values clustered by gene, with samples ordered by condition and replicate and annotated accordingly. (C) Comparison TPM of *LMNA*, *ZMPSTE24*. (D) Clustering of individual replicates of proteomics per line after PCA. (E) Comparison expression (log2) Lamin A/C and ZMPSTE24 from proteomics data (F) MA plot showing interesting up- or downregulated proteins (CTRL vs LKO (0/0)). (G) MA plot showing interesting up- or downregulated proteins (CTRL vs ZKO (0/0)).
